# Sequenceserver: a modern graphical user interface for custom BLAST databases

**DOI:** 10.1101/033142

**Authors:** Anurag Priyam, Ben J. Woodcroft, Vivek Rai, Alekhya Munagala, Ismail Moghul, Filip Ter, Mark Anthony Gibbins, HongKee Moon, Guy Leonard, Wolfgang Rumpf, Yannick Wurm

## Abstract

The dramatic drop in DNA sequencing costs has created many opportunities for novel biological research. These opportunities largely rest upon the ability to effectively compare newly obtained and previously known sequences. This is commonly done with BLAST, yet using BLAST directly on new datasets requires substantial technical skills or helpful colleagues. Furthermore, graphical interfaces for BLAST are challenging to install and largely mimic underlying computational processes rather than work patterns of researchers.

We combined a user-centric design philosophy with sustainable software development approaches to create Sequenceserver (http://sequenceserver.com), a modern graphical user interface for BLAST. Sequenceserver substantially increases the efficiency of researchers working with sequence data. This is due to innovations at three levels. First, our software can be installed and used on custom datasets extremely rapidly for personal and shared applications. Second, based on analysis of user input and simple algorithms, Sequenceserver reduces the amount of decisions the user must make, provides interactive visual feedback, and prevents common potential errors that would otherwise cause erroneous results. Finally, Sequenceserver provides multiple highly visual and text-based output options that mirror the requirements and work patterns of researchers. Together, these features greatly facilitate BLAST analysis and interpretation and thus substantially enhance researcher productivity.

## Introduction

The dramatic drop in cost of DNA sequencing since 2007 (http://genome.gov/sequencingcosts) has led to a rapid increase in the number of researchers working with unpublished sequence data from newly obtained transcriptomes, genomes and metagenomes. This has created many opportunities for novel biological research in fields including medicine (Qin *et al*., 2010), developmental biology (Santos *et al*., 2015), microbiology (Hugenholtz & Tyson, 2008), ecology and evolution (Ellegren, 2014; Seehausen *et al*., 2014). Pursuing such opportunities often involves querying newly obtained sequence data with specific aims such as determining the presence or absence of a sequence in a specific sample, obtaining sequence to enable molecular cloning, quantitative PCR or phylogenetic analyses, inferring the names or functions of putative genes, and identifying sequence and structural variation within and between species.

At a basic level such analyses involve comparing one or more query sequences with one or more databases each containing between thousands and hundreds of millions of sequences. This is typically performed with BLAST (Basic Local Alignment Search Tool; Altschul *et al*., 1990, 1997; Camacho *et al*., 2009), likely the most commonly used bioinformatics tool (>100,000 citations; see also Van Noorden *et al*., 2014; Korf *et al*., 2003). For each query sequence, BLAST heuristically identifies database sequences that are similar and outputs relevant comparisons (Camacho *et al*., 2009). To perform BLAST searches, biologists typically rely on graphical web interfaces provided by large organizations such as the National Center for Biotechnology Information (NCBI; Johnson *et al*., 2008) or the European Bioinformatics Institute (Goujon *et al*., 2010). These are valuable resources for published data, but despite efforts to encourage and facilitate early data deposition (Kaye *et al*., 2009) it can take months or even years before new sequence data is available on such central repositories.

How can researchers perform BLAST queries on new or unpublished data? First, BLAST software can be freely downloaded. However, using it directly requires substantial command-line and Unix skills (http://www.ncbi.nlm.nih.gov/books/NBK52640/) which many biologists lack (Smith, 2013). Furthermore, BLAST outputs are text-based; they lack potentially helpful graphical visualization. Second, queries and interpretation can be outsourced to bioinformatician collaborators or technicians with appropriate skills. Finally, the most appropriate solution for many laboratories would be to have access to a shared graphical user interface to run BLAST on private datasets. Several free and open source software packages make this possible, including ViroBLAST (Deng *et al*., 2007), GMOD’s BlastGraphic (O’Connor *et al*., 2008), NCBI’s wwwblast (Tao, 2006), NCBI’s Amazon Cloud BLAST (NCBI, 2014), and Galaxy BLAST (Cock *et al*., 2015). Unfortunately, setting up such interfaces is challenging: they are either deprecated and no longer maintained, or have complex installation dependencies such as custom configuration of the Apache web server (Fielding & Kaiser, 1997) or CGI paths (Gundavaram, 1996). Commercial software packages overcome some such problems, but their cost makes them inaccessible to many laboratories.

Once set up, a further challenge is using the graphical user interface. For example, users typically must choose between multiple BLAST algorithms (TBLASTX, TBLASTN, BLASTX, BLASTN, BLASTP) before entering query data or choosing search databases (e.g., Tao, 2006; Deng *et al*., 2007; O’Connor *et al*., 2008; NCBI, 2014; Cock *et al*., 2015; Johnson *et al*., 2008). This implicitly requires detailed knowledge of BLAST algorithms and available databases - yet even experienced BLAST users make frequent mistakes and must revisit their decisions. This design could be improved because only a single algorithm is appropriate for most combinations of input sequence and database. Many such inadequacies of BLAST user interfaces exist. This is likely because there is only a limited culture of user interface and user experience design for scientific software (Macaulay *et al*., 2009) including for bioinformatics (Pavelin *et al*., 2012).

Here we present Sequenceserver, a free software package designed to help overcome the largest of the aforementioned hurdles. By combining user-centric design (Garrett, 2011; Javahery *et al*., 2004; Pavelin *et al*., 2012) with modern software development and visualization approaches, Sequenceserver aims to increase the productivity of bioinformaticians setting up web servers for running BLAST on custom datasets, and for biologists performing and interpreting BLAST searches.

## Results

Sequenceserver is a graphical user interface wrapper for BLAST designed with extensive consideration of user experience and user interface (Garrett, 2011; Javahery *et al*., 2004; Pavelin *et al*., 2012). Below we provide overviews of Sequenceserver’s major features that facilitate setting up a BLAST server, performing queries and interpreting results.

### Assisted installation and configuration

Sequenceserver can be rapidly installed and configured by biologists with only little Unix experience for individual use or for shared use as part of a research group. Entering sudo gem install sequenceserver into a terminal will download and install Sequenceserver on standard Unix operating systems including Mac and Linux. Once installed, entering sequenceserver into a terminal downloads NCBI BLAST+ (Camacho *et al*., 2009) if necessary and prompts the user to indicate the location of a directory containing database sequences in the standard FASTA format or in BLAST’s specific BLASTDB database format. Sequenceserver then recursively scans the directory for FASTA files that have not yet been formatted as BLASTDBs. For each such FASTA file, Sequenceserver detects whether the FASTA file contains nucleotide or amino acid sequences and determines a suitable name to display the corresponding BLASTDB in the search form. Sequenceserver then confirms with the user if the FASTA file should be formatted as BLASTDB. Here, the user can modify the proposed name and optionally enter a relevant taxonomic identifier. The location of the directory containing database sequences is stored in a configuration file for subsequent use. Finally, Sequenceserver launches its built in web server (Harris & Haase, 2012) making the graphical user interface (Figure 1 and below) accessible from a web browser locally at http://localhost:4567 or over the network at http://user-ip:4567; it automatically shows all available BLASTDBs, segregated according to nucleotide vs. amino acid sequence type.

**Figure 1.**
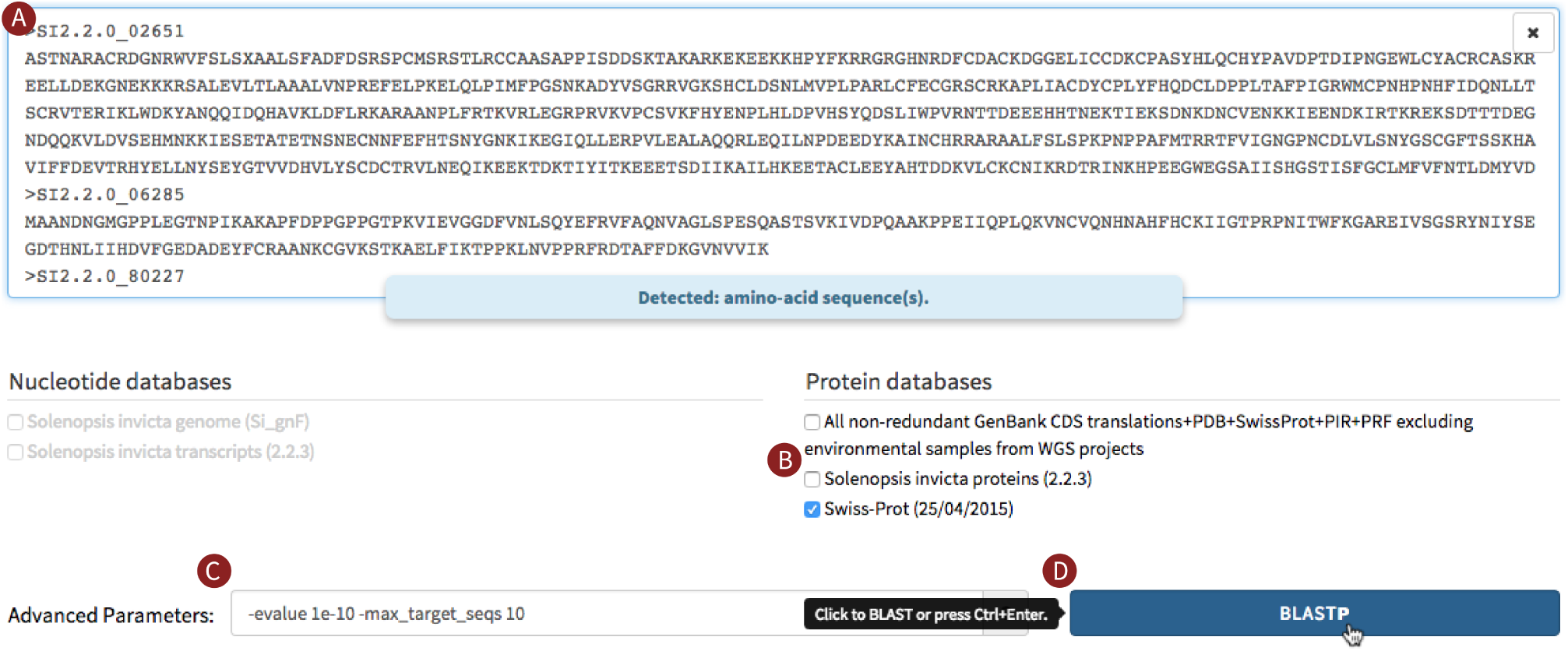
Partial screenshot of the query interface. Dark red letters highlight the steps involved and some specific features. **A:** Three or more sequences were pasted into the query field (typewriter font; only the identifier is visible for the third sequence); a message confirms to the user that these are amino acid sequences. **B:** The Swiss-Prot protein database was the first database to be selected. As a result, additional database selections are limited to protein databases; nucleotide databases are disabled. **C:** The user entered (optional) advanced parameters to constrain the results to the 10 strongest hits with evalues stronger than 10^−10^. **D:** The BLAST button is automatically activated and labeled “BLASTP” as this is the only possible basic BLAST algorithm for the given query-database combination. As the user’s mouse pointer hovers over the BLASTP button, a tooltip indicates that a keyboard shortcut exists for this button.

Additional configuration possibilities include setting the number of threads to be used by BLAST, integrating Sequenceserver with web servers such as Nginx (Reese, 2008) or Apache (Fielding & Kaiser, 1997), adding password protection, customizing the interface colors and layout, and adding custom links from hits to external resources such as genome browsers (see online documentation).

### Assistive BLAST query submission interface

Sequenceserver uses techniques including analysis of user input, interactive feedback and simple algorithms to streamline BLAST query submission and to prevent common errors that can cause BLAST to fail or generate misleading results. The user first types, pastes or drag-and-drops one or multiple FASTA format query sequences into a text-field (Figure 1). In the unlikely event that the user combined nucleotide sequences and amino acid sequences, an alert message is shown and the BLAST button will remain disabled to avoid BLAST generating meaningless results. Subsequently, the user selects one or several BLAST databases using checkboxes. Once a first database is selected, additional database selections are limited to those of the same type (i.e., either nucleotide or protein) to eliminate the risk of users combining incompatible databases that would cause BLAST to fail. Once a valid query is entered and a database is selected, the BLAST submission button activates. For most query-database combinations, the single possible basic BLAST algorithm will be used (Supplementary Figure S1). When multiple algorithms are appropriate (e.g., nucleotide query and nucleotide database: BLASTN and TBLASTX are both appropriate), a pull-down in the BLAST submission button allows the user to toggle between them. Sequenceserver’s automatic algorithm selection reduces the risk of attempting to perform impossible BLAST queries. Finally, Sequenceserver includes an “advanced parameters” field providing access to all standard BLAST parameters available in the command-line (Camacho *et al*., 2009).

### Display of BLAST results

As a result of performing a BLAST query, an HTML report including graphical overviews is shown in the web browser (Figure 2; an interactive version of this figure is at http://sequenceserver.com/paper/fig2interactive). This report will feel familiar to users of NCBI BLAST but includes many additions and revisions for improved navigation, interpretation and follow-up analysis. For example, the report can be downloaded in HTML, XML and tab-delimited table formats; similarly a FASTA file containing all hit sequences can be downloaded. The HTML report clearly brings out the structure of results: If a user submitted multiple query sequences, a clickable index of queries is shown on one side. All queries, hits and BLAST HSPs (high-scoring segment pairs) are numbered to facilitate navigation. For each query, identified hits are summarized in a table and an overview graphic. This graphic indicates the strength and locations of alignment of each hit (hits with stronger e-values are darker).

**Figure 2.**
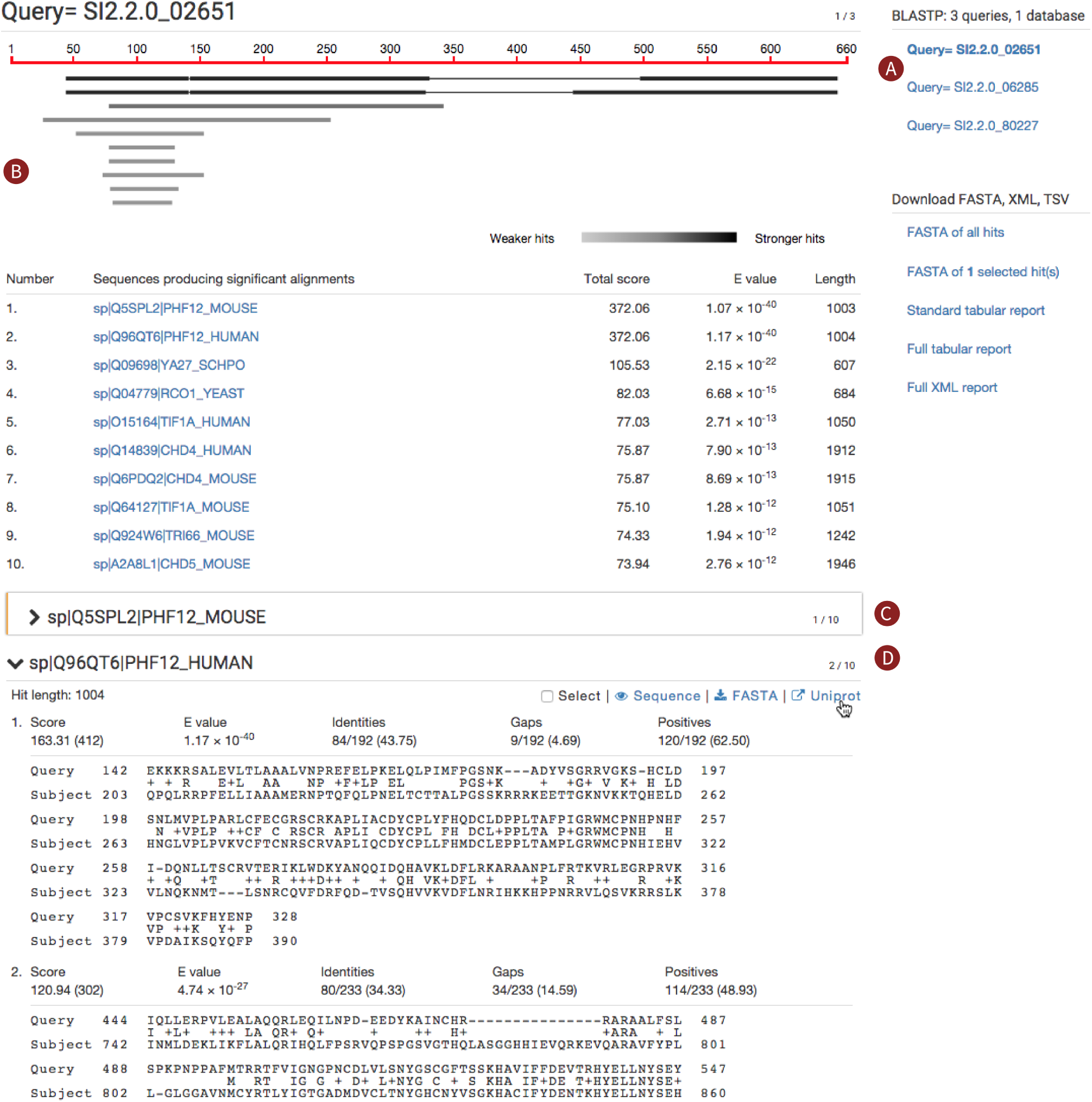
Screenshot of a Sequenceserver BLAST report. *An interactive version of this figure is at* http://sequenceserver.com/paper/fig2interactive. Three amino acid sequences were compared against the Swiss-Prot database using BLASTP with an evalue cutoff of 10^−10^ and keeping only the 10 strongest hits per query. This screenshot shows a portion of the results for the first query. Dark red letters highlight some of the specific features of this report. **A:** An index overview summarizes the query and database information and provides clickable links to query-specific results. **B:** Results for the first query are shown. These include a graphical overview indicating which parts of the query sequence aligns to each hit, a tabular summary of all hits, and alignment details for each hit. **C:** The first hit is selected for download; its alignment details have been folded away. **D:** The user is studying the second hit; the mouse pointer hovers over the link to the hit’s UniProt page.

Each hit includes multiple links for subsequent analysis. For example, each hit to a sequence with an NCBI or UniProt identifier includes a link to the relevant page; links to private genome browsers or other sites can be added (see online documentation). Additionally, each hit includes one link to download the full sequence in FASTA format and another link to display the sequence in the browser. This sequence viewing interface (Supplementary Figure S2) includes GenBank-style visualization for readability and displays appropriate coordinate information when the mouse pointer hovers over one or selects multiple residues (Gómez *et al*., 2013). Furthermore, each hit includes a checkbox making it possible to simultaneously download a selection of multiple hit sequences as a single FASTA file.

## Discussion

We created Sequenceserver to overcome many of the challenges of performing BLAST on custom datasets. Below we review known applications of Sequenceserver, compare it to alternatives, consider its compatibility with tools for follow-up analyses and discuss future directions.

### Applications and relevance

Sequenceserver has accelerated our own research (Gotzek *et al*., 2011; Ingram *et al*., 2012; Wurm *et al*., 2011; Wang *et al*., 2013; Kulmuni *et al*., 2013; Privman *et al*., 2013; Mondav *et al*., 2014; Schrader *et al*., 2014; Nygaard & Wurm, 2015) and that of others. Indeed, despite this being the first formal publication reporting Sequenceserver, the software has already been cited for research on emerging model organisms (Wurm, 2015) such as sea cucumber (Rowe *et al*., 2014), starfish (Elphick *et al*., 2013; Semmens *et al*., 2013, 2015), falcons (Seim *et al*., 2015), the sugar-apple tree (Gupta *et al*., 2015), the Hessian fly *Mayetiola destructor* (Shreve *et al*., 2013) and the house plant *Streptocarpus rexii* (Chiara *et al*., 2013), as well as for research in bioadhesion (Rodrigues *et al*., 2014), environmental microbiology (Castro *et al*., 2015), and general research on BLAST (Sharma & Mantri, 2014).

Google Analytics referral links to http://sequenceserver.com, social web statistics, and community engagement through the mailing list indicates growing adoption of Sequenceserver (Table 1). For example, Sequenceserver is among the 10% most frequently downloaded packages on the biogems portal for bioinformatics software (Bonnal *et al*., 2012, > 28,000 downloads), and is a main querying mechanism for community genomic databases including the SymGRASS database of sugarcane orthologous genes (Belarmino *et al*., 2013), the *Drosophila suzukii* genome database (Chiu *et al*., 2013), the ant genomes database (Wurm *et al*., 2009), the planarian database (Brandl *et al*., 2015), the butterfly genomics database (http://blast.lepbase.org), a portal for exploring low-complexity amino acid sequences (Kirmitzoglou & Promponas, 2015), the *Amborella* database (http://amborella.huck.psu.edu/), a *Fusarium* database (http://www.fusariumnrpspks.dk/), two ash genome projects (https://geefu.oadb.tsl.ac.uk/ and http://www.ashgenome.org/) and an echinoderm transcriptome database (http://echinodb.uncc.edu/blast/). We are aware of at least 50 publicly accessible Sequenceserver instances, but precise usage statistics are difficult to confirm, as there is no centralized list of deployments. Our experience with deployment suggests that public servers are in the minority with most Sequenceserver instances being created for use by individual researchers or small groups as needed on the personal computers of researchers. For instance, the authors collectively run 12 private servers and only 4 public servers at the time of this writing.

**Table 1.** Usage and impact statistics of Sequenceserver at the time of writing.

| Metric | Statistic |
| --- | --- |
| Hits to sequenceserver.com in the last 12 months | 11,050 |
| Mailing list users | 102 |
| Mailing list threads | 137 |
| RubyGems downloads | 29,500 |
| GitHub stars | 80 |
| Twitter mentions | 102 |
| Sites with publicly available instances | 52 |

Sequenceserver is also used as an educational resource (e.g., http://culture-bioinformatics.org/blast.shtml) to provide entry-level bioinformatics students a controlled environment to run BLAST queries and explore the effects of changing BLAST parameters on the results. Indeed, SequenceServer allows the students to access their classroom’s BLAST server from anywhere and the instructor to control the software and databases versions so that the results for any given query are consistent and predictable, as is critical for a classroom setting and not feasible using publicly-available servers such as NCBI.

### Alternatives and compatibility with tools for follow-up analysis

Multiple alternatives to Sequenceserver similarly provide graphical user interfaces (GUIs) for BLAST (Tao, 2006; Deng *et al*., 2007; O’Connor *et al*., 2008; Cock *et al*., 2015; NCBI, 2014). However, they have three broad levels of shortcomings (summarized in Supplementary Table S1). First, most of these tools first require other software to be installed and include little or no automation of common tasks such as downloading and configuring BLAST+ software or adding custom databases. Installation of such tools is out of reach of most biologists. In contrast, biologists with only limited Unix experience are able to set up Sequenceserver. Importantly, this ease of setup does not limit possible usage options. Indeed, experienced power users retain the flexibility of deploying Sequenceserver on personal computers, compute clusters and the cloud, customizing all aspects of Sequenceserver, and migrating from its built-in webserver to industry-strength standards.

Second, the query submission interface of alternatives to Sequenceserver are designed around how BLAST is implemented rather than accompanying the user through their thought and work process. For example, in some cases (Cock *et al*., 2015; Johnson *et al*., 2008) users must select a BLAST algorithm even before they have entered the query sequence and selected the databases to search. This is counterintuitive as a user’s thought process generally begins with their query sequence. In other cases (Tao, 2006; Deng *et al*., 2007; O’Connor *et al*., 2008; NCBI, 2014), users must select a BLAST algorithm after they have entered the query or after they have selected the databases to search. This is more in-line with users workflow, but both approaches fail to consider that only a single basic BLAST algorithm is applicable for most query-database combinations, thus the users wastes time choosing the appropriate algorithm. Furthermore, none of the alternative GUIs consider that the user may erroneously select an algorithm that is inappropriate for their query-database combination, or consider that the user may erroneously combine nucleotide and amino acid sequence in a single query. In both scenarios, BLAST will either fail or incorrectly interpret the nucleotides adenine (A), thymine (T), cytosine (C) and guanine (G) as the amino acids arginine, threonine, cysteine and glutamine.

Finally, most GUIs (Tao, 2006; Deng *et al*., 2007; O’Connor *et al*., 2008; Cock *et al*., 2015; NCBI, 2014) either present the default HTML output of command-line BLAST or slightly reformat it. They thus lack many of the features of Sequenceserver that facilitate navigation, interpretation and follow-up analyses including exporting analysis results in different formats. Sequenceserver uses an approach that allows exporting a single analysis in many different formats. As part of this, Sequenceserver’s HTML report is generated from scratch, making it much easier to extend it with additional features.

Several commercial providers offer alternatives to Sequenceserver (e.g., BlastStation, Geneious, CLC). While such tools can overcome some of shortcomings listed here, they have other drawbacks. These include costs, risks in terms of reproducibility and long-term sustainability, limited flexibility for sharing with distant colleagues, and limited customization possibilities.

BLAST reports from Sequenceserver (e.g., in XML and tab-delimited formats) can be used as input for analysis tools that work directly with BLAST output. Thus users can access an instance of Sequenceserver running on a high-performance computer to create a BLAST report which they subsequently use with tools that facilitate specific analyses [e.g., MEGAN (Huson *et al*., 2007), Blast2GO (Conesa *et al*., 2005), BLAST2GENE (Suyama *et al*., 2004), EPoS (Griebel *et al*., 2008)], provide different visualization facilities [e.g., Kablammo (Wintersinger & Wasmuth, 2015), BLASTGrabber (Neumann *et al*., 2014), Circoletto (Darzentas, 2010), JAMBLAST (Lagnel *et al*., 2009), BOV (Gollapudi *et al*., 2008)] or facilitate importing BLAST results into a spreadsheet application or an SQL database [e.g., MuSeqBox (Xing & Brendel, 2001), Zerg (Paquola *et al*., 2003), Batch BLAST extractor (Pirooznia *et al*., 2008), BioParser (Catanho *et al*., 2006)]. A comparison of tools to interpret and work with BLAST output is provided by Neumann *et al*. (2013).

### Future directions

The community of Sequenceserver developers will continue adding novel functionalities to our software in a public and open manner. We expect improvements along three main lines. First, additional output visualization facilities will be immensely beneficial for making sense of large datasets (Nielsen *et al*., 2010). In particular, this will involve integrating visualization tools developed by others (Gollapudi *et al*., 2008; Neumann *et al*., 2013; Gómez *et al*., 2013; Wintersinger & Wasmuth, 2015; http://wurmlab.github.io/tools/genevalidator/), while HMMER provides high-sensitivity for comparisons against sequence profiles (Eddy, 2009). Similarly, DIAMOND is up to 20,000 times faster than BLASTX or BLASTP; this can be useful when datasets are particularly large (Buchfink *et al*., 2015, e.g., metagenomics). New versions of Sequenceserver and their features will be announced on Sequenceserver’s mailing list and Github repository.

### Conclusion

Recent improvements in DNA sequencing technologies are enabling ever smaller groups of biologists to create previously unimaginable datasets on diverse ranges of species (Koboldt *et al*., 2015; Ellegren, 2014). Realizing that a major challenge for many biologist researchers is to effectively query such datasets, we built the Sequenceserver graphical frontend for BLAST using modern user interface and user experience paradigms. Our software is particularly enabling for researchers and communities focusing on species for which genomic data is novel as it empowers them to independently and relatively easily perform BLAST queries and share datasets. At the base of this software are modern software development technologies and approaches that have over the last few years become heavily used by the small groups of software developers at the core of many internet startup companies (Ries, 2011; Thiel & Masters, 2014). We are excited that the computational tools underlying basic scientific research can benefit from such technical possibilities (Altschul *et al*., 2013) and appropriate consideration of user experience (Pavelin *et al*., 2012). The increased usability, maintainability and sustainability of well-built tools ultimately translate into increased agility and productivity of researchers and in turn the impacts of their breakthroughs.

## Methods

### Programming environment

We developed Sequenceserver from scratch rather than basing our work on the NCBI’s initial Perl/CGI www-blast wrapper (Tao, 2006) to reduce technical debt (Lehman, 1980). The core of Sequenceserver is written in the Ruby programming language (Flanagan & Matsumoto, 2008) popular for creating dynamic websites (Ruby & Thomas, 2011) and bioinformatics tools (Goto *et al*., 2010), while JavaScript and HTML/CSS are used for layout and interactions in the web browser. We use multiple preexisting tools and libraries to facilitate development: The lightweight framework Sinatra (Harris & Haase, 2012) is used to create URL endpoints to load search form and run a BLAST search from the browser. BLAST searches are delegated to the compiled command line version of BLAST (Camacho *et al*., 2009); we use Ox (https://github.com/ohler55/ox) to parse BLAST XML and create the HTML report. Underscore (http://underscorejs.org/), HTML5 Shiv (https://github.com/afarkas/html5shiv), jQuery (http://jquery.com), jQuery UI (http://jqueryui.com), Web-shim (https://afarkas.github.io/webshim/demos/), and Twitter Bootstrap (http://getbootstrap.com) libraries are used to create a uniform scripting environment (for dynamic aspects of the user interface) and a consistent look-and-feel (for visual layout) across most browsers. The d3 (http://d3js.org/) and BioJS (Gómez *et al*., 2013) libraries are used respectively for generating the graphical overview and the sequence viewing interface.

### Sustainable software development approach

Throughout the development of Sequenceserver we followed six modern software engineering practices designed to facilitate and accelerate development while improving the long-term sustainability of the software (Prlić & Procter, 2012; Wilson *et al*., 2014; http://www.software.ac.uk/resources/guides-everything). First, we used an open source and agile development approach (Shore & Warden, 2007) involving frequent incremental improvements, peer review and frequent deployment on our own servers and within the community. Second, we structured the software according to object-oriented paradigm (Weisfeld, 2008), thus leading to a clean separation of different parts of the code. Third, we followed two important software development principles: “don’t repeat yourself” (DRY) leads to fewer lines of code and thus fewer bugs, and makes it easier to read and understand code than if similar commands are repeated in several places (Hunt & Thomas, 2000); “keep it simple, stupid” (KISS) reduces unnecessary complexity and thus lowers risks and leads to higher maintainability (Raymond, 2003). Fourth, we reuse widely established software packages and libraries (see programming environment) to avoid reimplementation. This accelerates our work and reduces the amount of Sequenceserver-specific code, which in turn further reduces the likelihood of adding bugs (Sametinger, 1997). Fifth, we developed unit tests (Ammann & Offutt, 2008) and integration tests (Ammann & Offutt, 2008) for many parts of Sequenceserver’s code, and used continuous integration (https://travis-ci.org/) to ensure these tests are automatically run whenever any change is made to the code, thus increasing the chances of rapidly detecting errors. Sixth, we use the rubocop code analyser (https://github.com/bbatsov/rubocop) to ensure that our code respects the Ruby community style guide in terms of names of variables and methods, code structure and code formatting. Such respect of style standards makes code more accessible to other programmers and scientists than if code is inconsistently styled or if we had chosen our own conventions (Martin, 2008; Wurm, 2015). Finally, we use the Codeclimate service (http://codeclimate.com) for automatic detection of potential problems with software design.

### Graphical user interface design principles

To ensure a fluid user experience that increases researcher productivity, we designed Sequenceserver around eight modern user interface design principles. First, the interface contains only essential information so as to minimize distractions for the user. Second, the information is laid out in a clear and hierarchically structured manner. As part of this, we paid special attention to typography, using Roboto (https://www.google.com/fonts/specimen/Roboto) for headings and Open Sans (https://www.google.com/fonts/specimen/Open+Sans) for normal text. These free, contemporary typefaces were designed to maximize legibility and overall aesthetics across electronic devices and print media. Third, we used automation where possible to minimize the number of decisions required from the user. For example, based on query type and databases selection we limit the choices for algorithm selection (except in the case of nucleotide-nucleotide search only a single BLAST algorithm is possible; see Supplementary Figure S1). Fourth, we use interactive visual feedback and cues for step-by-step discovery of the workflow. For example, the BLAST button remains disabled until the user has provided query sequence(s) and selected target databases. If the user tries to click the BLAST button while it is disabled a tooltip indicates that a required input is missing. Similarly, selection of protein databases is automatically disabled if the user has already selected a nucleotide database (and vice versa). Fifth, we remain consistent and contextual with regards to user interaction. For example, notification of sequence type does not depend on how query sequence was provided. This notification is shown below the query sequence input field - where the user’s eyes are likely to go after query input - instead of using a global designated notification area or displaying pop-up windows that can be disruptive or are ignored. Similarly, a “clear query” button is shown only after the user has provided query sequence(s) and is positioned where a user is likely to look for it. Sixth, we try not to let the advantages of a graphical interface and efforts to create an easily accessible user experience limit the scope of what the user can do. For example, all possible advanced BLAST search options can be entered via a generic input field. Similarly, tooltips over report download links are only shown after the mouse pointer has hovered for at least 500ms. This delay means most users won’t be bothered by tooltips after they have used the interface a few times. Seventh, we exploit intuitive human notions of colors. For example, if the user erroneously tries to combine nucleotide and amino acid sequences in the query, the query input-area is gently highlighted using a red border to indicate an error. At a different level, in the graphical overview shown for each query, the color of each hit indicates its strength, with stronger e-values being darker. Finally, the wording of error messages is similar to informal human conversation to create empathy and familiarity, which may also clarify that Sequenceserver is built by a community of scientists. For example, if a user encounters a bug in Sequenceserver, the error popup reads “Oops! Something went wonky. Below is the error detail. Please could you report it to our Google Group.”

### Community building

We took four measures to help create a community and allow easy customization of our software. First, we performed all development in an open manner on the GitHub source code sharing/development site (http://github.com). This ensures that bioinformatician users can easily access the source code and customize the software. Indeed, such customization can be easily merged back into the main codebase of the software; several GitHub users are included as coauthors of this manuscript because of such contributions. Second, we provide extensive documentation as part of the source code and on the website http://sequenceserver.com. Third, we created a dialog with users (via GitHub issue tracking as well as a specific Google Groups mailing list); here users ask and answer each other’s questions and help each other regarding issues arising with customization, installation and specific datasets. Finally, we isolated the most commonly used customization code into separate files (a configuration file and a specific file for adding links from BLAST search results to custom databases such as custom genome browsers or other databases). Isolating the parts of code that a bioinformatician is most likely to edit facilitates customization and reduces risks when upgrading the software.

## Acknowledgments

The authors would like to thank the large community of Sequenceserver users and contributors for their feedback and input throughout this project.

## Supplementary Information

**Figure S1.**
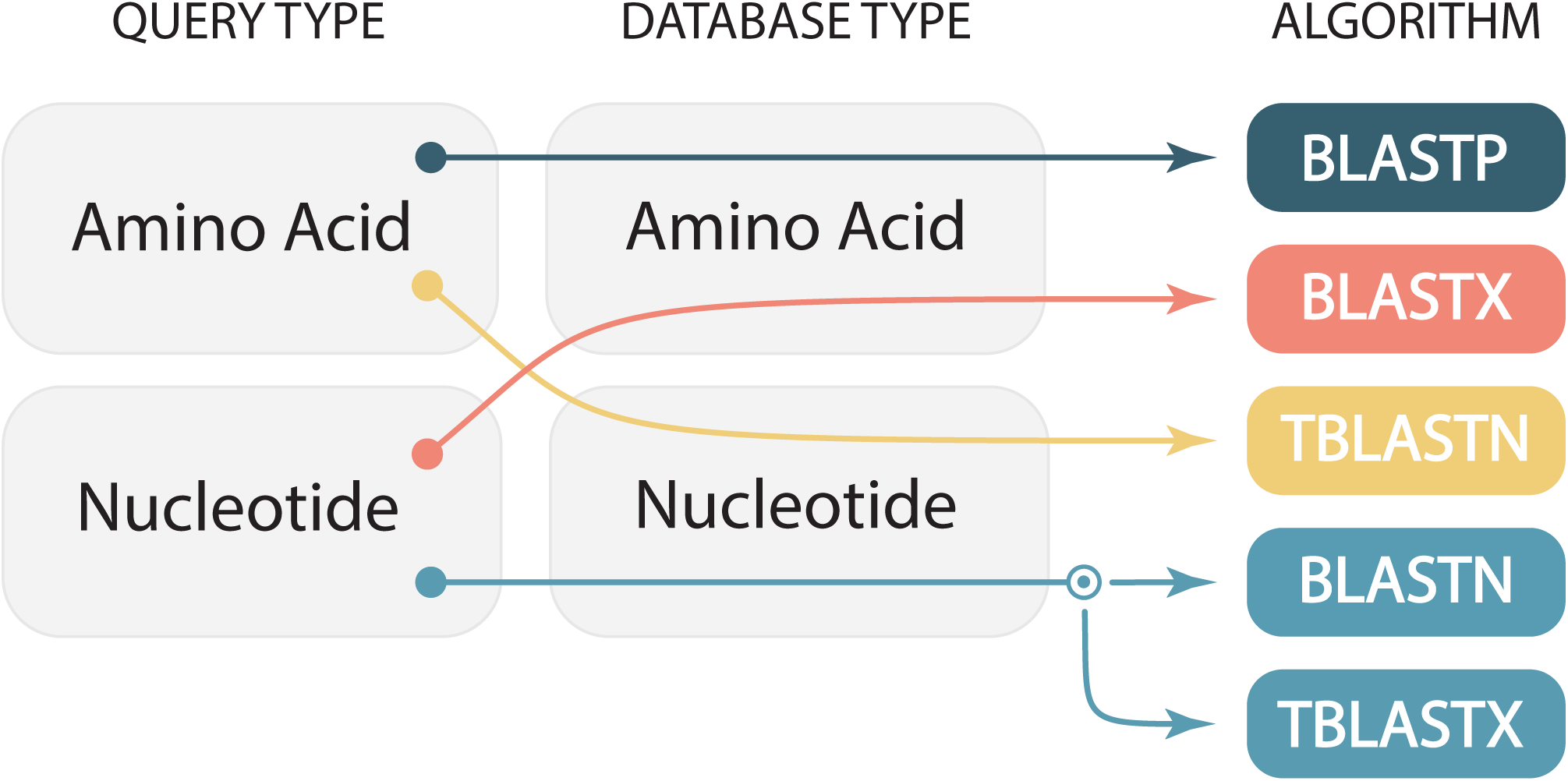
Automatic BLAST algorithm selection. BLAST includes five basic algorithms (right column). Arrows indicate how Sequenceserver automatically selects an appropriate BLAST algorithm based on the sequence types of query (left column) and selected databases (middle column). For the first three combinations of query and database types, only one algorithm is possible. The circle indicates that for nucleotide query and nucleotide database, the user can choose between BLASTN and TBLASTX.

**Figure S2.**
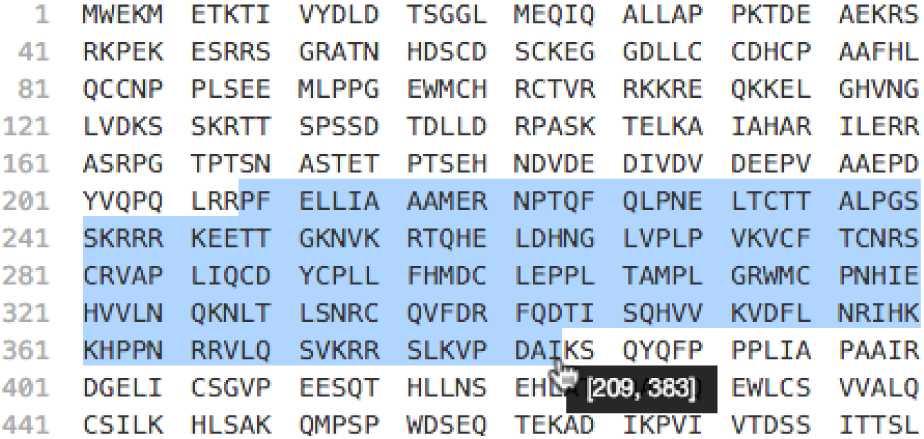
Screenshot of viewing interface for a protein hit sequence. Residues are grouped in multiples of 5, and the coordinates of the first residue of each line are shown in gray. Here, the user selected a range of amino acids; their specific coordinates are shown in a tooltip. The viewing interface for nucleotides is similar.

**Table S1.** Alternatives to Sequenceserver. Presence or absence of features in BLAST GUIs is highlighted across four broad categories: Installation and configuration, BLAST query submission, Output, and Development status. Preferred features are indicated in green.

|  |  | ViroBlast | NCBI's<br>wwwblast | GMOD's<br>BlastGraphic | Galaxy Blast | NCBI Amazon<br>Cloud BLAST | Sequenceserver |
| --- | --- | --- | --- | --- | --- | --- | --- |
| Reference and/or URL |  | Deng et al 2007 | Tao T (2006)<br>wwwblast: Setup<br>and usage. http://<br>www.ncbi.nlm.nih.gov/staff/tao/<br>URLAPI/<br>wwwblast/. | Oconnor et al<br>2008 | Cock et al 2015 | NCBI (2014)<br>BLAST searches<br>at the cloud.<br>http://<br>blast.ncbi.nlm.nih.gov/blast/docs/<br>BLAST_at_AWS.<br>pdf. | This publication |
| Installation and configuration | Specific software requirements | Apache web<br>server, PHP | Apache web<br>server, CGI | Apache web<br>server, Perl and<br>several Perl<br>modules from<br>CPAN | Galaxy server | Only runs on<br>Amazon Web<br>Services | Ruby<br>programming<br>language<sup>1</sup> |
|  | BLAST software download | Manual | Manual | Manual | Automatic<sup>2</sup> | Included | Automatic |
|  | BLAST configuration | Manual | Manual | Manual | Automatic | Included | Automatic |
|  | Detection and formatting of new<br>datasets | Manual | Manual | Manual | Manual | Manual | Automatic/<br>assisted |
|  | Adding new datasets to interface | Manual<br>configuration | Manual<br>configuration | Manual<br>configuration | Manual<br>configuration | Manual<br>configuration | Automatic |
|  | Support for NCBI's current BLAST+<br>software (Camacho et al 2009). | Yes | No | Yes | Yes | Yes | Yes |
|  | Updating software while preserving<br>configuration | Involved | Involved | Involved | Involved | Involved | Trivial |
|  | Can be integrated with standard<br>web servers for community<br>website | Yes | Yes | Yes | Yes | Yes | Yes |
| BLAST query submission | Automatic BLAST algorithm<br>selection | No | No | No | No | No | Yes |
|  | Prevent user from mixing protein<br>and nucleotide query sequences | No | No | No | No | No | Yes |
|  | Prevent user from running BLAST<br>with impossible query-database-<br>algorithm combinations | No | No | No | No | No | Yes |
|  | Ability to select multiple databases<br>for search | Yes | No | Yes | Yes | No | Yes |
|  | Ability to enter advanced options | Yes | Yes | Limited | Yes | Yes<sup>3</sup> | Yes |
| Output | Graphical overview of hits | No | Yes | Yes | No | No | Yes |
|  | Results clearly structured as in the<br>NCBI's public BLAST interface. | No | No | No | No | No | Yes |
|  | Ability to filter results | Yes: by e-value<br>and identity. | No | No | No | No | No |
|  | Link hits to custom external<br>resources | No | No | Yes | No | No | Yes |
|  | Ability to download hit sequences | Yes | No | No | No | No | Yes |
|  | Ability to download BLAST reports | No | Yes<sup>4</sup>: XML | No | No | Yes<sup>4,5</sup>: XML, Tab-delimited, JSON<br>& text | Yes: XML & Tab-delimited |
| Development status |  | Unclear | Inactive | Inactive | Active, open<br>community | Active,<br>developed only<br>by NCBI | Active, open<br>community |
| Notes |  | <sup>1</sup> Pre-installed on Mac. One-line installation through package managers on Linux.<br><sup>2</sup> Through 'ncbi_blast_plus' package from galaxy tool shed<br><sup>3</sup> Default: only e-value; configuration enables more options.<br><sup>4</sup> The format in which results are desired can be pre-selected from the search form.<br><sup>5</sup> Using the command line blast_formatter tool. |  |  |  |  |  |

